## supplemental_figure for "CamoTSS: analysis of alternative transcription start sites for cellular phenotypes and regulatory patterns from 5’ scRNA-seq data"

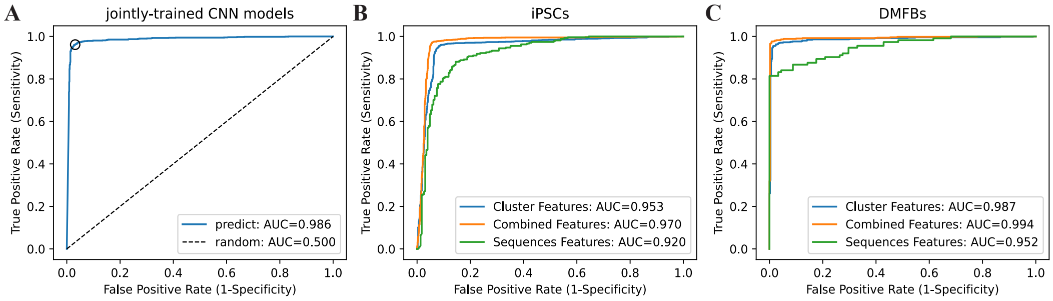


Figure S1: Prediction performance with different models, features and datasets. (A) Receiver operating characteristic curve for prediction of TSS clusters when combining reads-based (i.e., cluster features) and sequence-based features with joint training by concatenating the four reads-based features to the second last layer of CNN model. Shown is based on the combined dataset with iPSC and DMFB, same as main Fig. 1D. (B,C) ROC curves for iPSC (B) and DMFB (C) datasets with different feature groups by using logistic regression models. Panel B and C have the same model setting as main Fig. 1D.

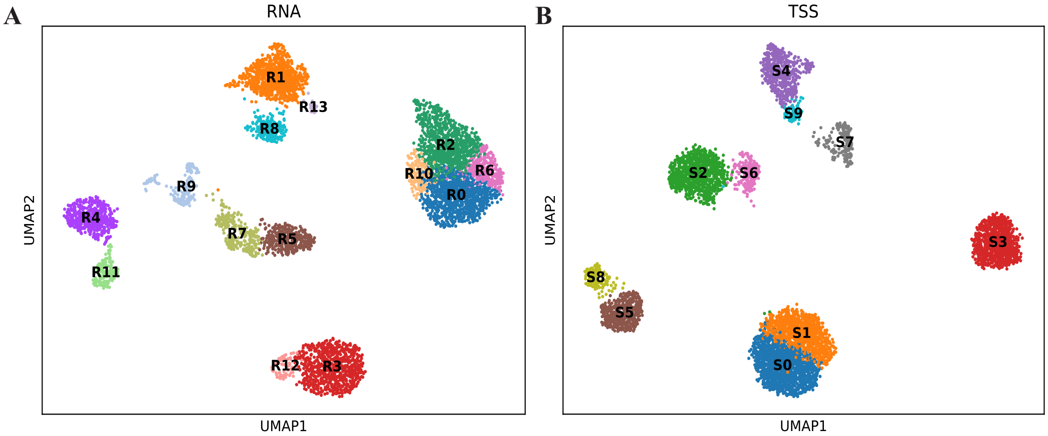


Figure S2: Uniform Manifold Approximation and Projection (UMAP) representation of 5732 single-cell transcriptomes clustered by gene expression (A) and TSS expression (B), colored by their according cell types. Prefix 'R' and 'S' denotes cell clusters of RNA and TSS, respectively.

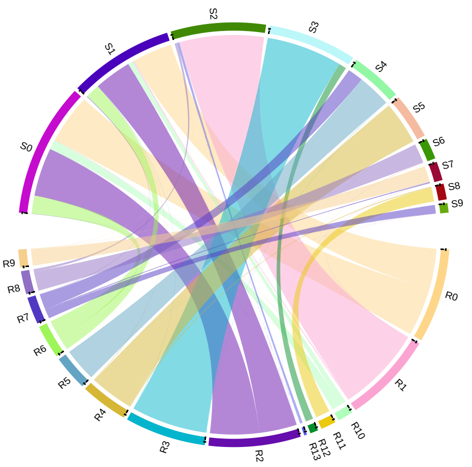


Figure S3: Chord diagram showing the relationship between all RNA-based (i.e., gene-level) clusters and all TSS-based clusters. Cluster IDs are the same as main Fig. 2A.

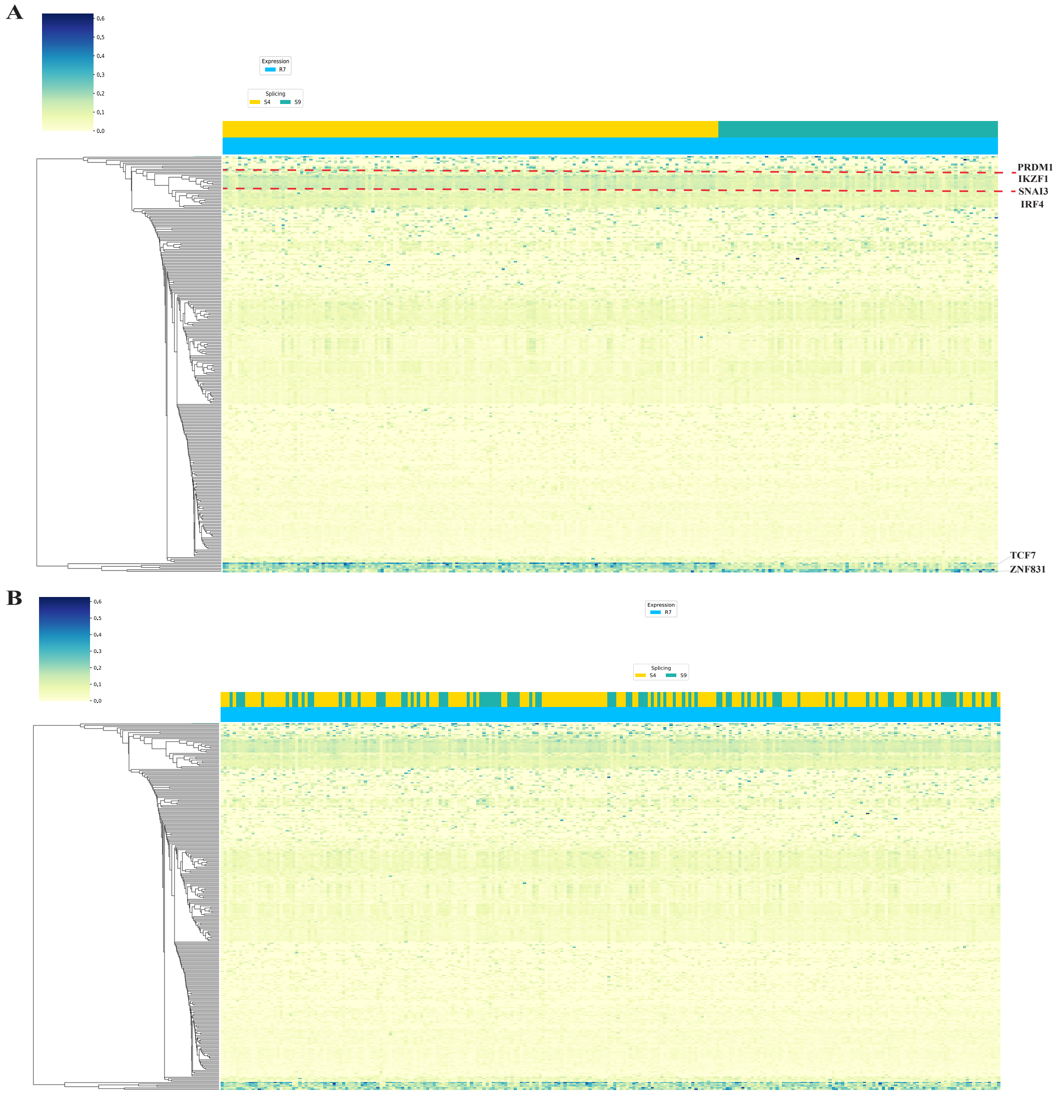


Figure S4: Heatmap showing SCENIC analysis of regulon (TF) activity in the R7 cluster of cells with S4 and S7 order (A) and random order (B).

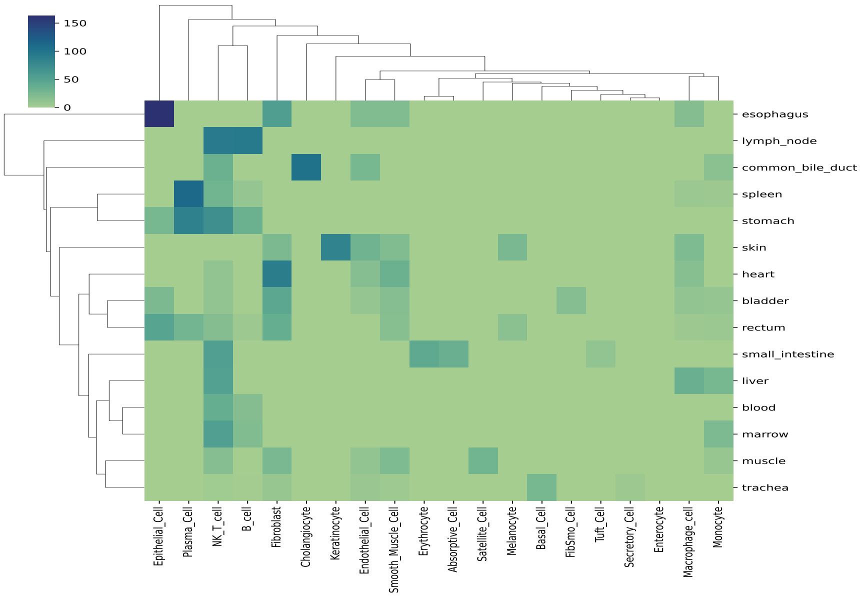


Figure S5: Heatmap showing the number of genes which have at least one TSS with a significantly differential ratio between one cell type vs all other cell types, across 15 organs

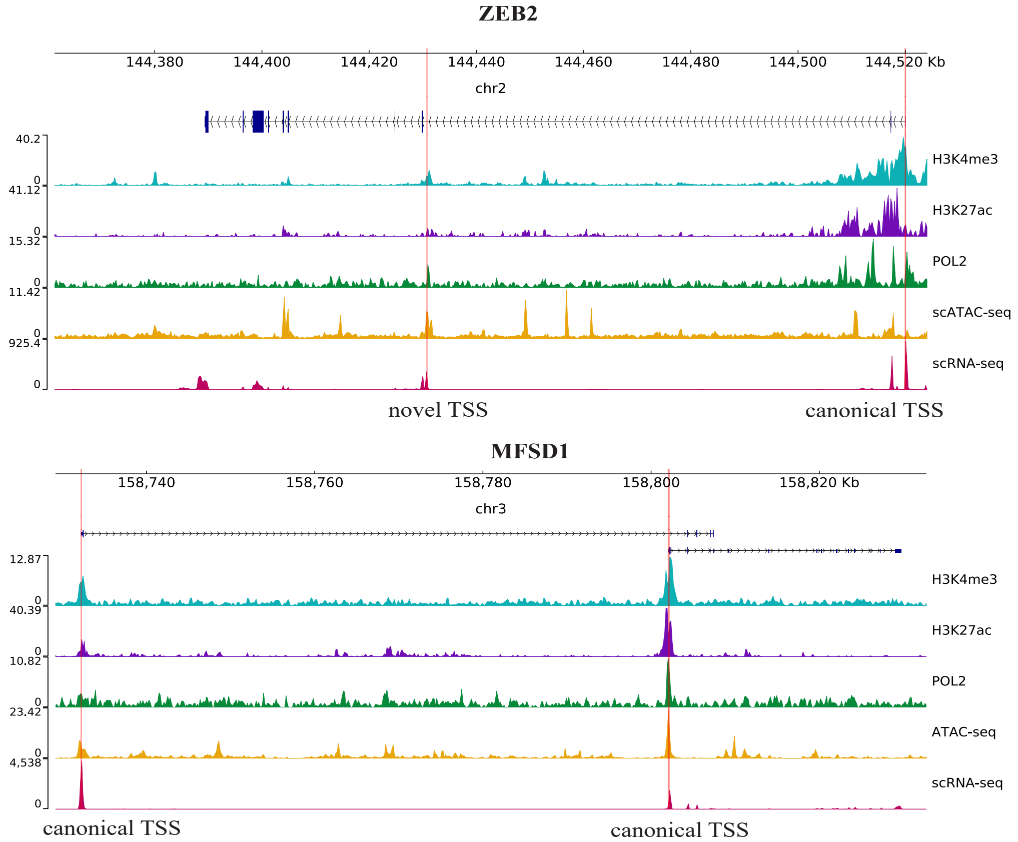


Figure S6: Tracks plot showing the coverage of histone modification mark (i.e H3K4me3 and H3K27ac), RNA POL2 and ATAC-seq of ZEB2 (Top) and MFSD1 (Bottom) in muscle and heart, respectively. The structure of genes is shown in UCSC style. The red line highlights the TSS detected by CamoTSS.

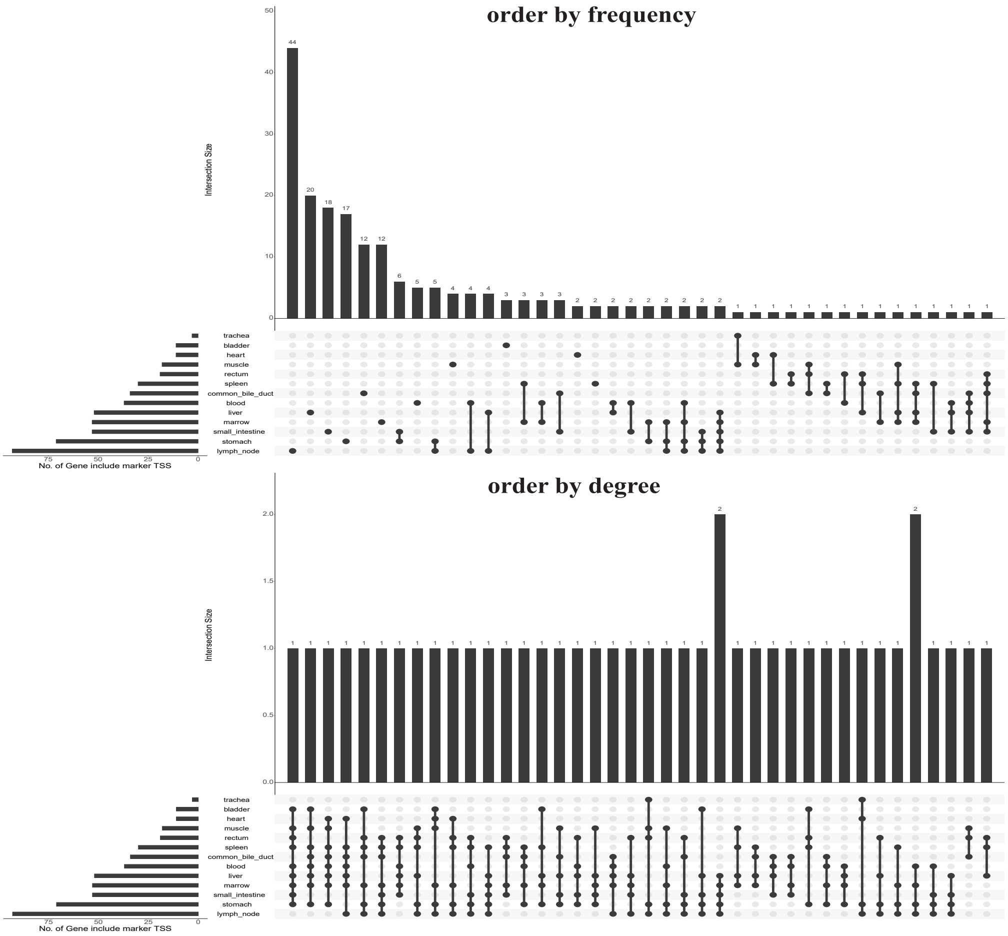


Figure S7: Upset plots displaying intersection of TSS marker in 13 organs. The upper panel is ordered by frequency. The bottom panel is ordered by the degree of intersection.

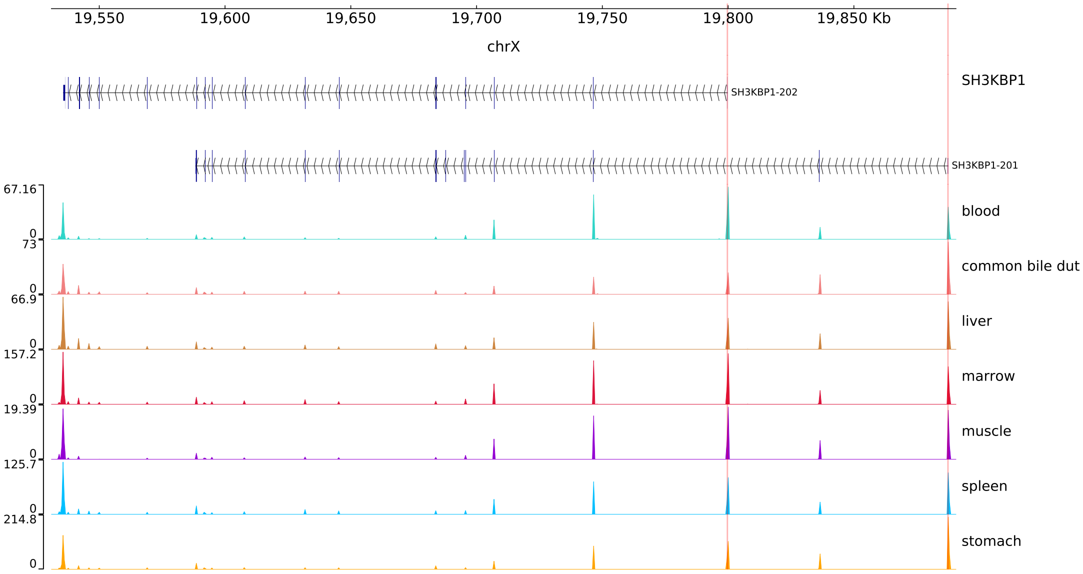


Figure S8: Tracks plot showing the coverage of SH3KBP1 in different organs in scRNA-seq data. Figure format is the same as Supp. Fig. S6.

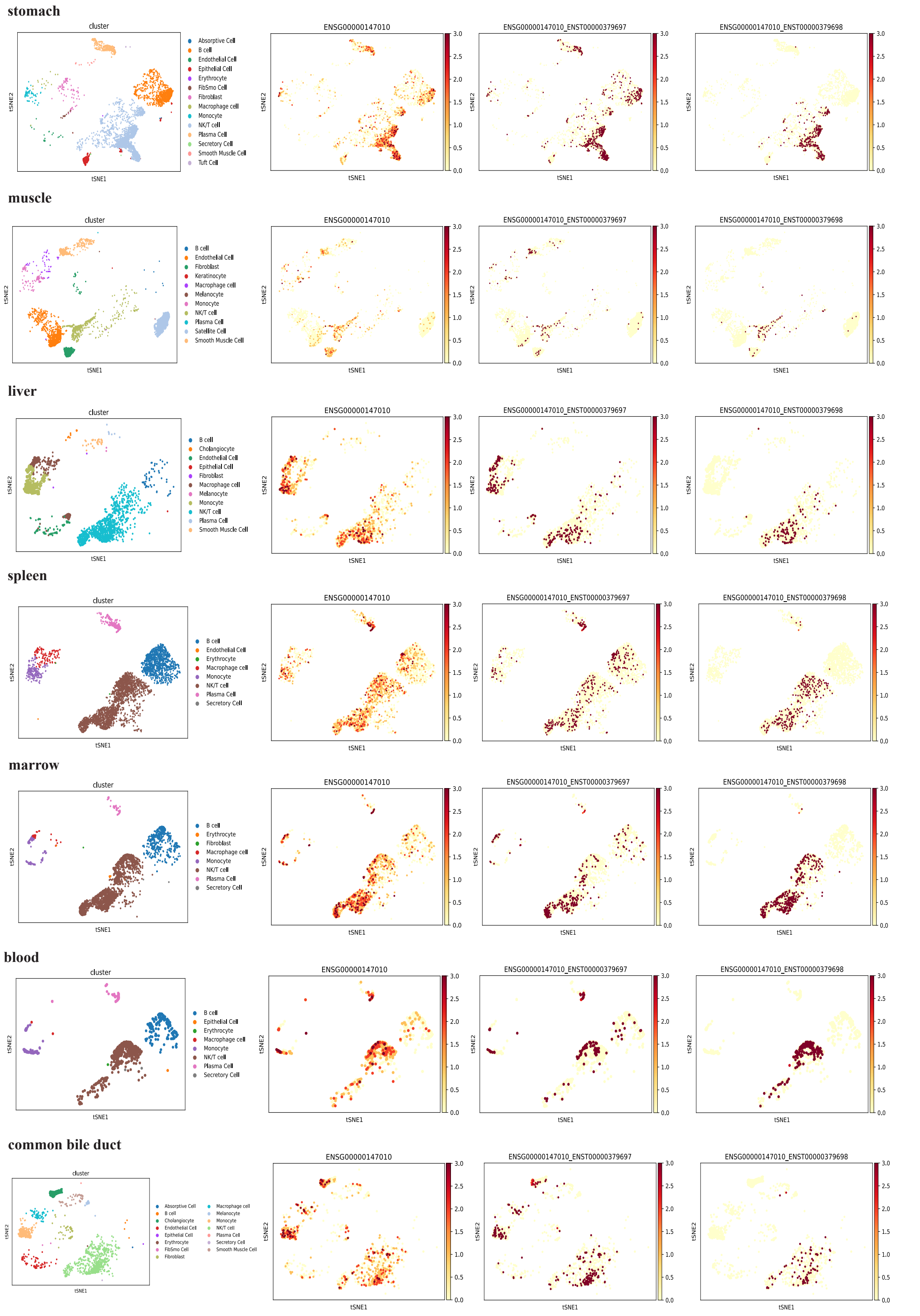


Figure S9: tSNE visualization showing cell clusters of different 7 organs (left panel). The SH3KBP1 gene-level expression (middle panel) and two SH3KBP1 TSSs (right panel) distribution in log1p(count) across cells superimposed on the t-SNE of the cells in 7 different organs.

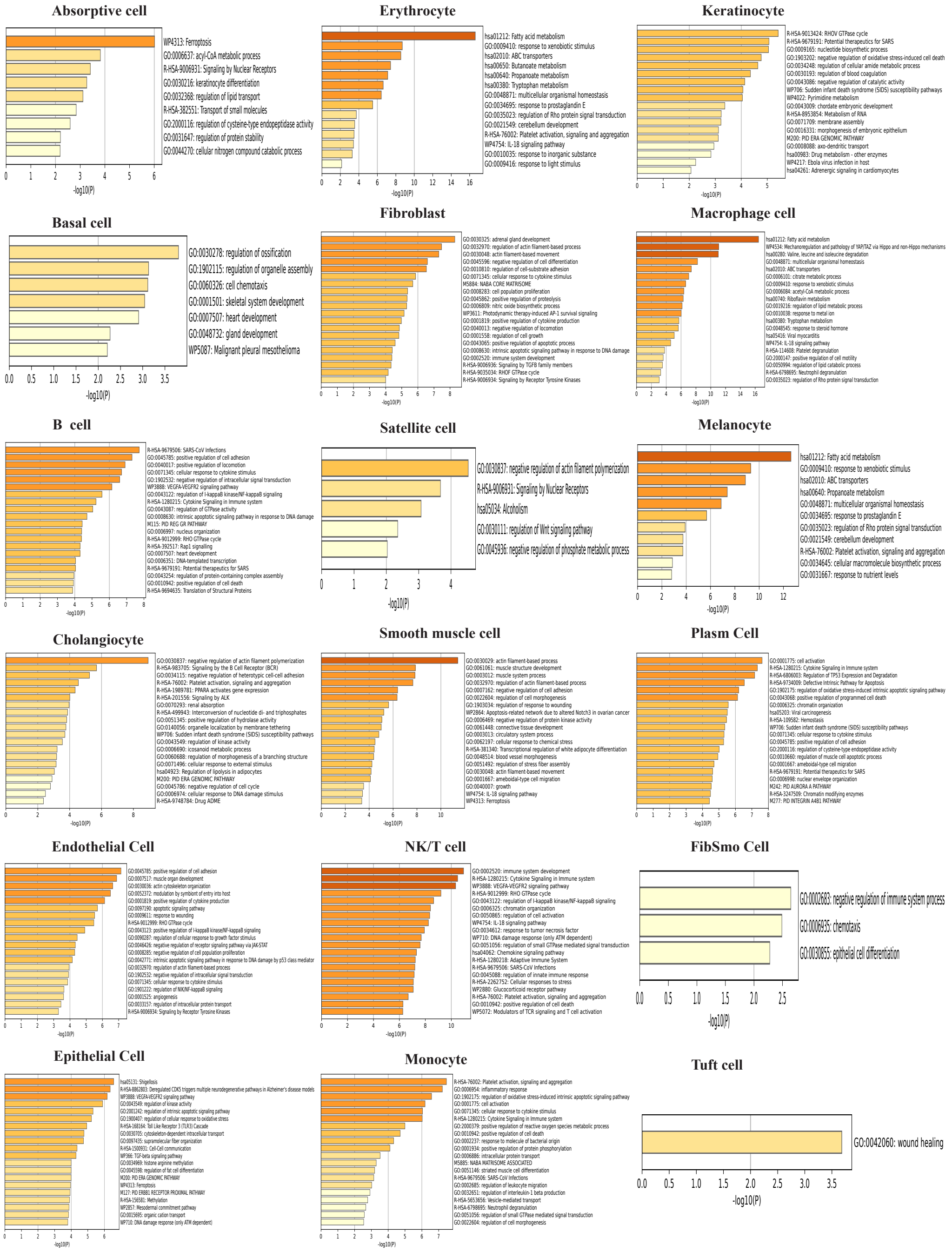

Figure S10: Bar plots of enriched GO terms over genes with significant alternative TSS usage.

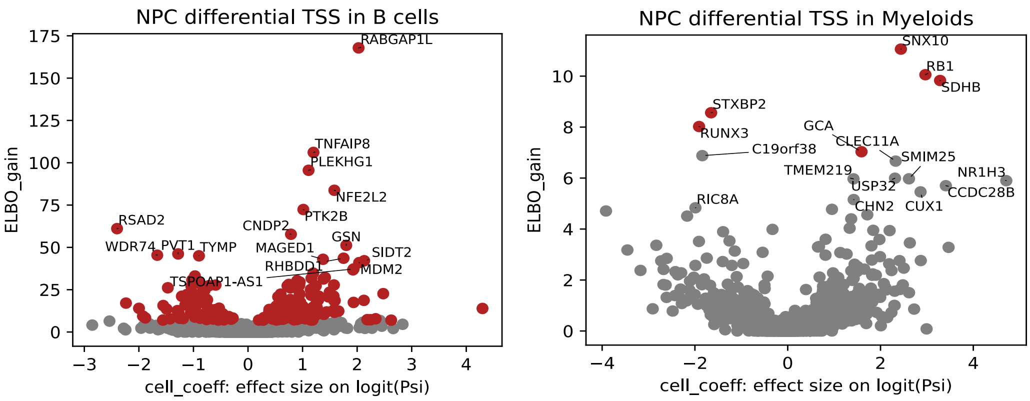


Figure S11: Volcano plots between ELBO_gain and effect size on logit(PSI) for detecting differential TSS usage between NLH and NPC patients, in B cells (left panel) and Myeloid cells (right panel). PSI value denotes the proportion of TSS1 between two major TSSs.

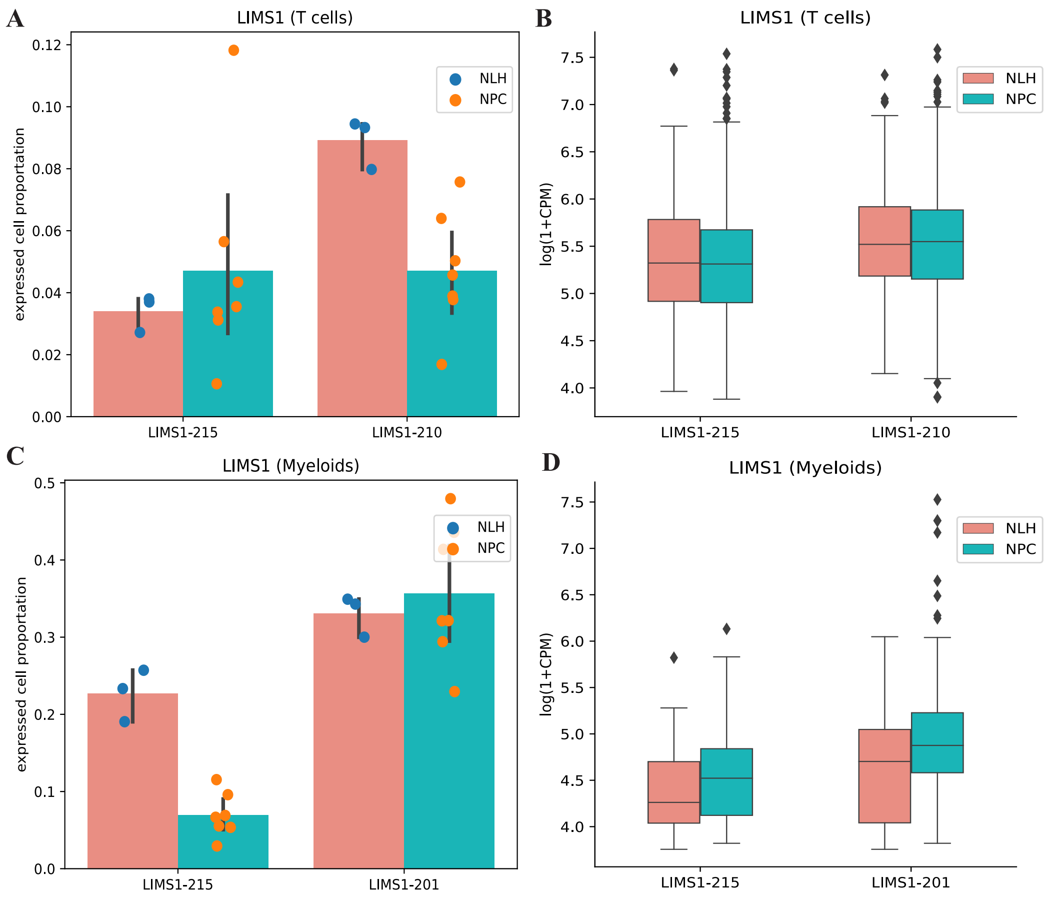


Figure S12: Expression profile of TSSs of LIMS1 in T cells and Myeloids. (A,C) Bar plot showing the expressed cell proportion of two alternative TSS of one gene between NLH and NPC patients, in T cells (A) and Myeloid cells (C). Box plots (B,D) showing the expression value of LIMS1 in the expressed cells in T (B) and Myeloid cells (D), corresponding to (A) and (C).

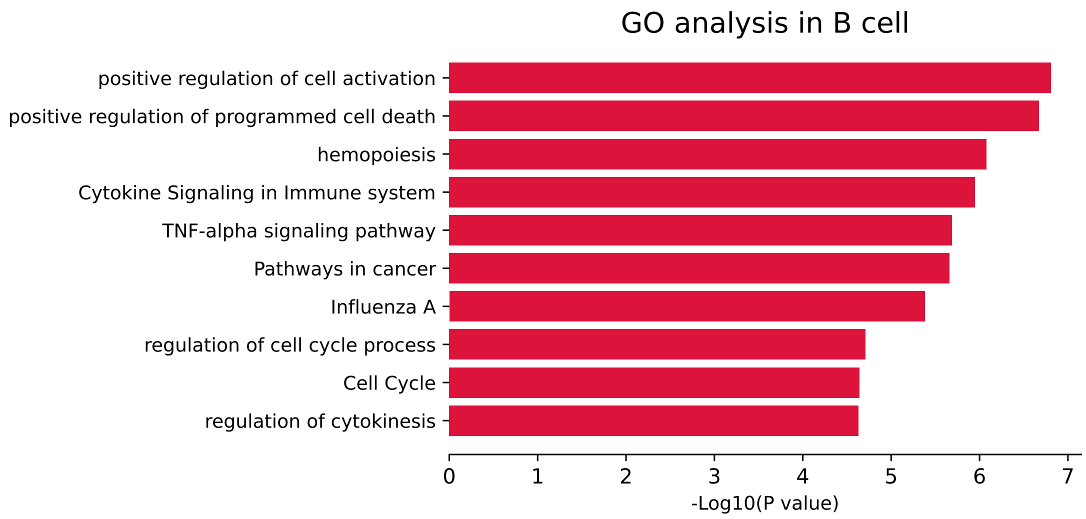


Figure S13: Bar plot showing the enriched terms of genes with differential TSS usage between NLH and NPC patients in B cells.

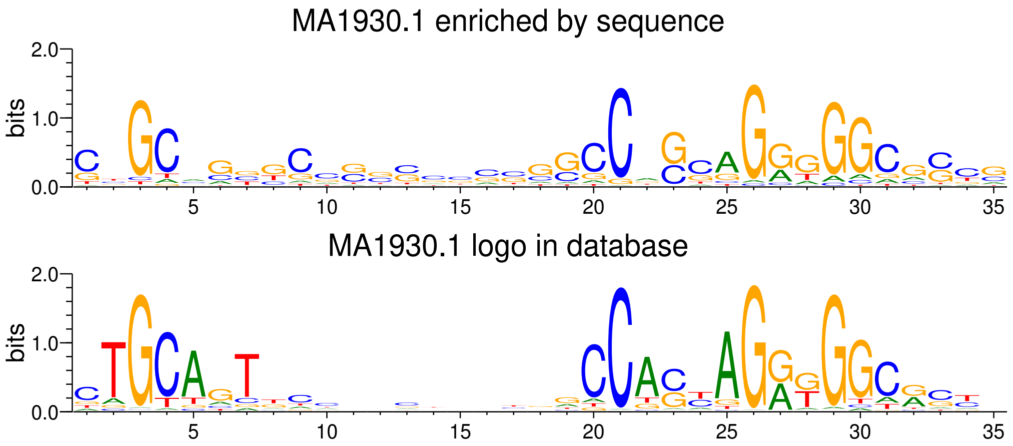


Figure S14: WebLogo of the base frequency of MA1930.1 (i.e. one motif of CTCF) enriched in the sequences detected by FIMO (top) and displayed in the JASPAR database (bottom).

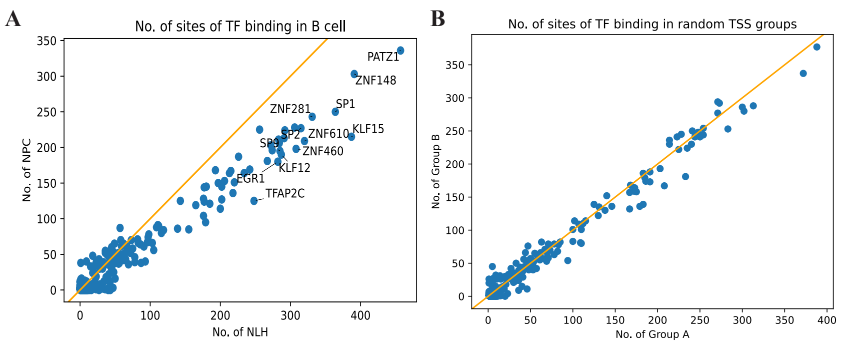


Figure S15: Binding frequency in different scenarios. (A) Scatter plot showing the binding frequency of human TFs detected by FIMO in B cells of NLH and NPC patients. (B) Scatter plot showing the binding frequency of human TF with two equal-sized random sequence sets of TSSs in NPC dataset.

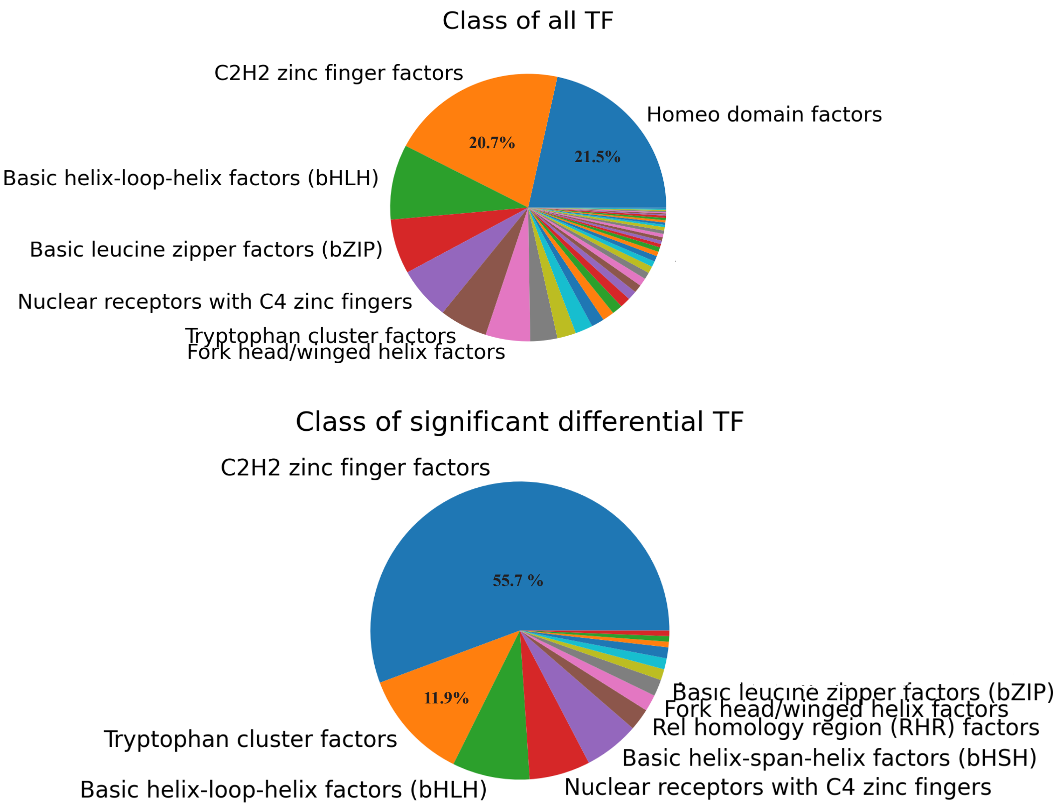


Figure S16: Pie chart showing the percentage of distinct classes of all TFs analyzed by FIMO (top panel). Pie chart showing the proportion of significant differential TFs detected by FIMO between NLH and NPC patients (Fisher's exact test).

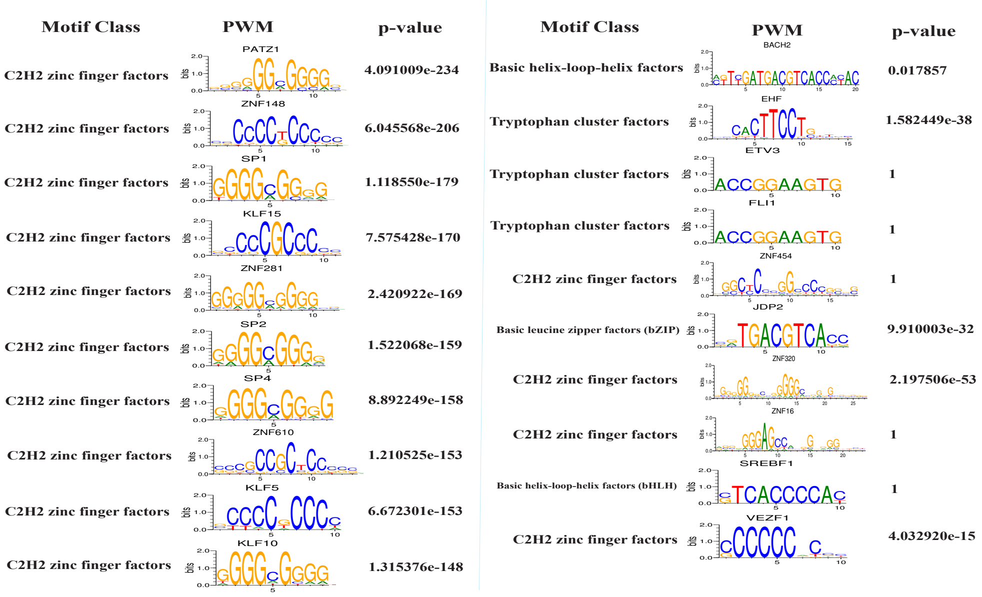


Figure S17: Weblogo showing the top 10 differential motifs and 10 randomly picked motifs. Strikingly, the 10 differential motifs are all from the C2H2 zinc finger class, while it's only occupies 40% in the 10 random motifs (close to the expectation from the background, shown in Fig. S16 top panel).

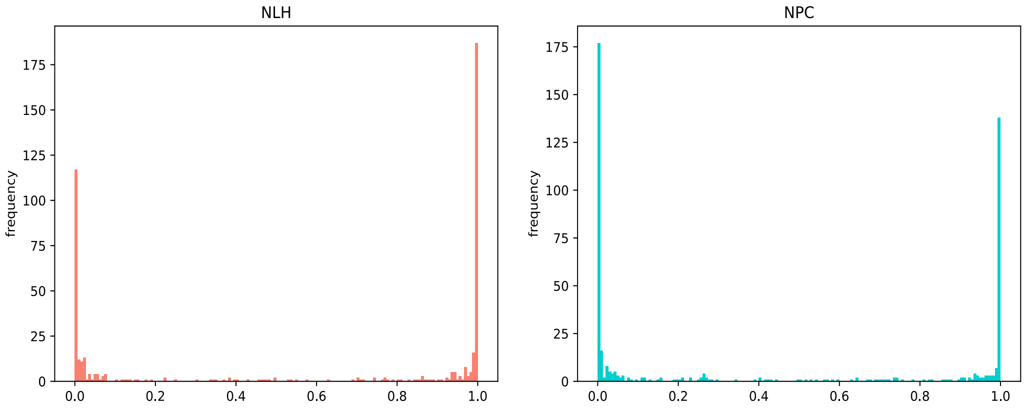


Figure S18: Histogram of probability of being positive samples predicted by our sequence-based CNN model, for the 528 TSS sequences elevated in NLH (left panel) and NPC (right panel).

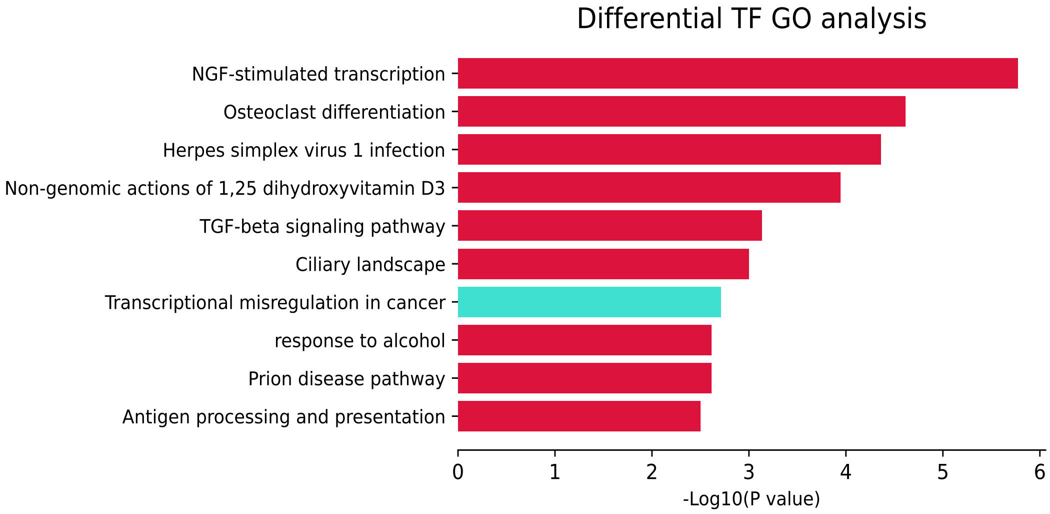


Figure S19: Bar plot showing the enriched GO terms of differential TFs with a distinct binding frequency between NLH and NPC patients. The blue bar highlights a highly related function with this dataset.

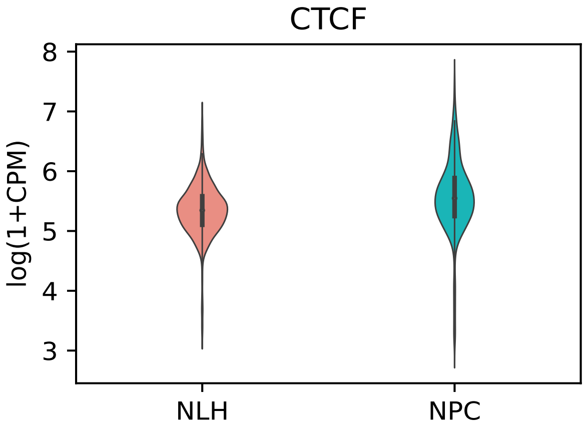


Figure S20: Violin plot showing the expression value of CTCF in the expressed cells between NLH and NPC patients.

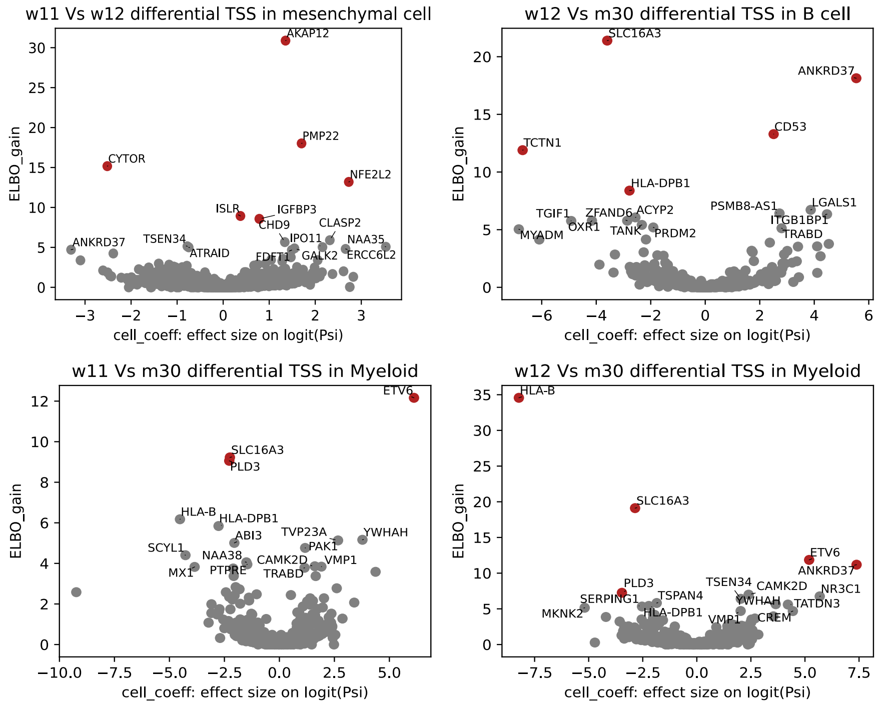


Figure S21: Volcano plots showing the relationship between ELBO_gain and effect size on logit(PSI) for all TSSs which can be detected by BRIE2. The PSI value denotes the proportion of TSS1 among the two major TSSs.

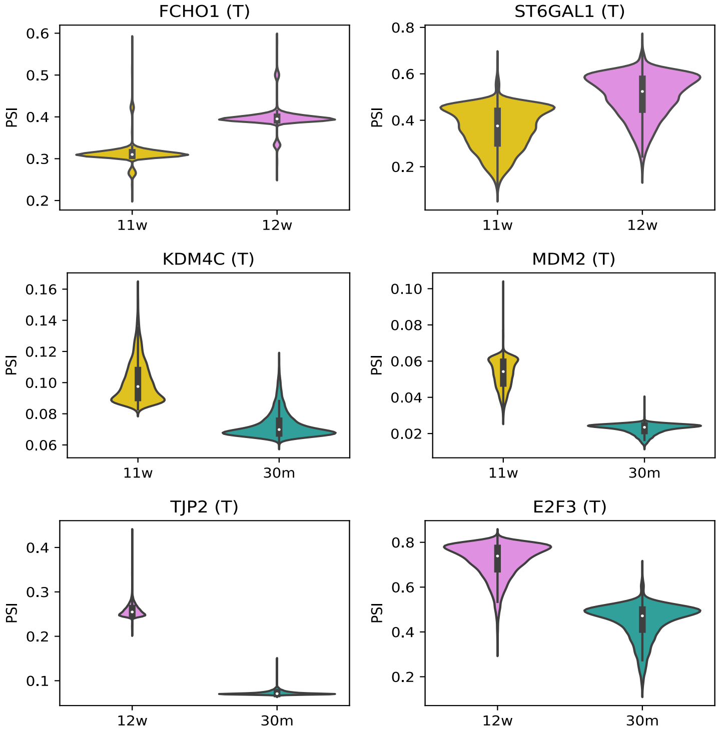


Figure S22: Violin plots of the significantly differential genes with alternative TSS usage but only between two development stages.

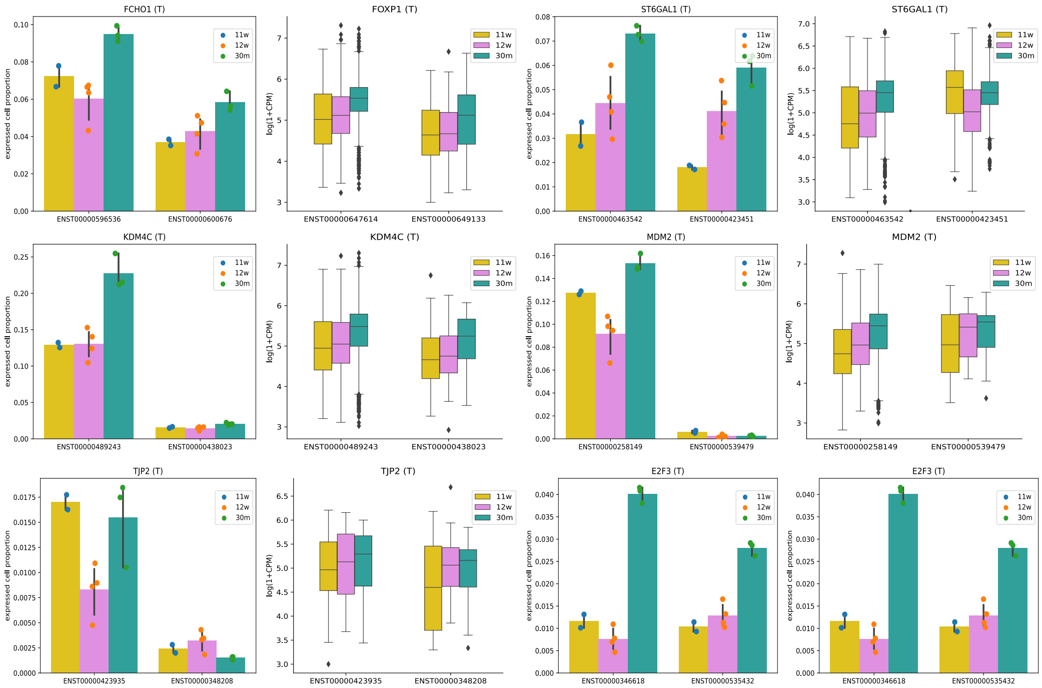

Figure S23: The expressed cell proportion and expression value of the example genes shown in Supp. Fig. S22.

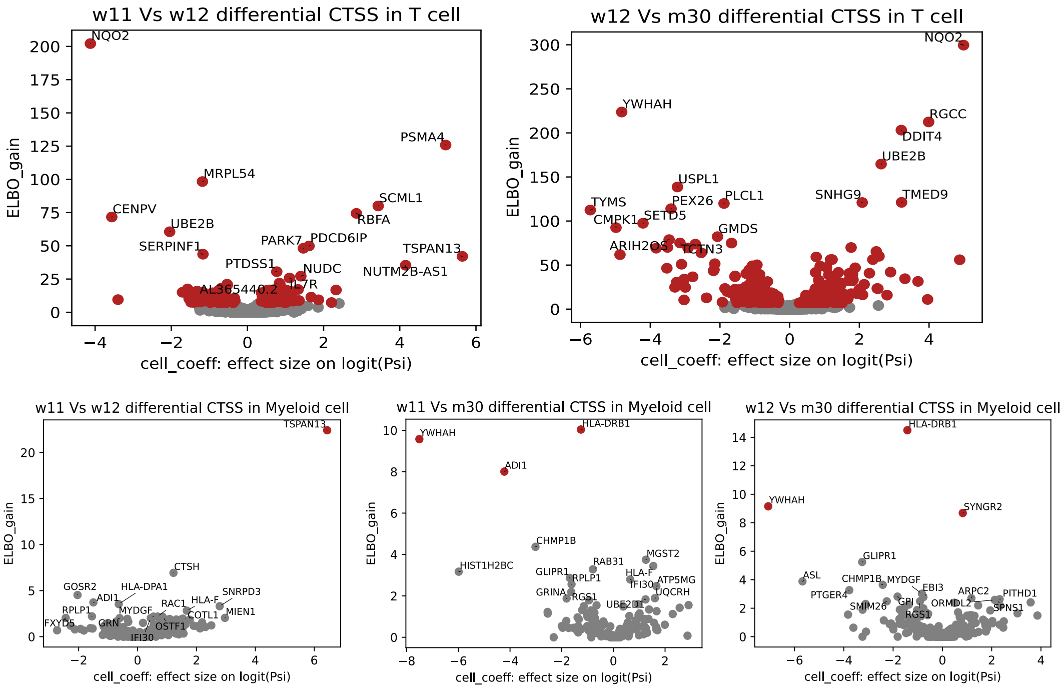

Figure S24: Volcano plots showing the relationship between ELBO_gain and effect size on logit(PSI) for differential CTSS between week11 and week12, week12 and month30 in T cells (Top) and in Myeloid cells (Bottom). The PSI value denotes the proportion of CTSS1 among the two major CTSSs.

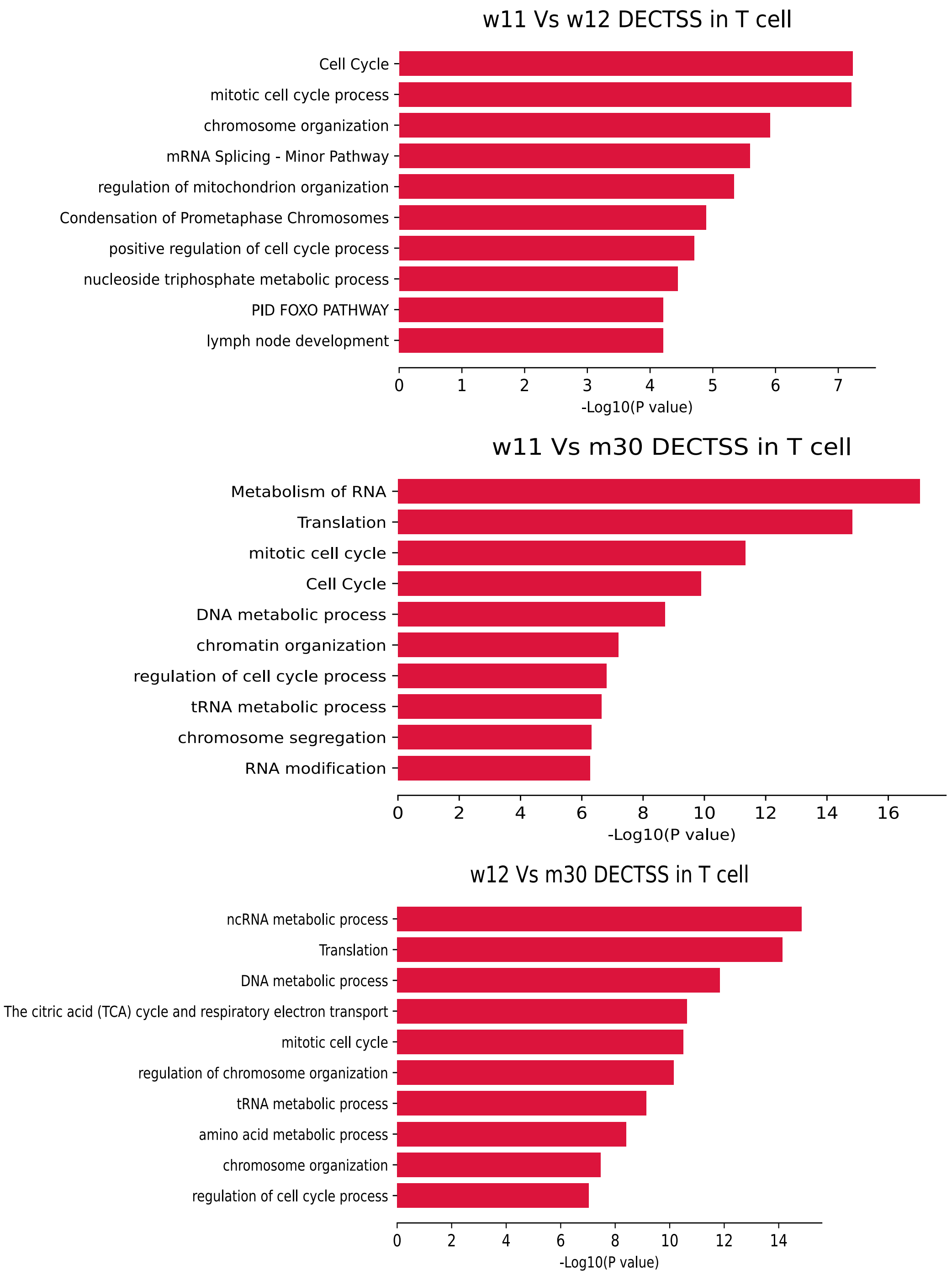


Figure S25: Bar graph of Gene Ontology enrichment analysis. Enrichment of gene ontology terms in gene list with a narrow shift within one TSS cluster respectively between 11-week and 12-week (top), 11-week and 30-month (middle), and 12-week and 30-month (bottom).
